# UPA-Seq: Prediction of Functional LncRNAs Using Differential Sensitivity to UV Crosslinking

**DOI:** 10.1101/332080

**Authors:** Taiwa Komatsu, Mari Mito, Koichi Fujii, Shintaro Iwasaki, Shinichi Nakagawa

## Abstract

While a large number of long noncoding RNAs (lncRNAs) are transcribed from the genome of higher eukaryotes, systematic prediction of their functionality has been challenging due to the lack of conserved sequence motifs or structures. Assuming that lncRNAs function as large ribonucleoprotein complexes and thus are easily crosslinked to proteins upon UV irradiation, we performed RNA-Seq analyses of RNAs recovered from the aqueous phase after UV irradiation and phenol-chloroform extraction (UPA-Seq). As expected, the numbers of UPA-Seq reads mapped to known functional lncRNAs were remarkably reduced upon UV irradiation. Comparison with ENCODE eCLIP data revealed that lncRNAs that exhibited greater decreases upon UV irradiation preferentially associated with proteins containing prion-like domains (PrLDs). Fluorescent in situ hybridization (FISH) analyses revealed the nuclear localization of novel functional lncRNA candidates, including one that accumulated at the site of transcription. We propose that UPA-Seq provides a useful tool for the selection of lncRNA candidates to be analyzed in depth in subsequent functional studies.

## Introduction

A large number of long noncoding RNAs (lncRNAs) are transcribed from the genomes of higher eukaryotes, and accumulating evidence suggests that lncRNAs are involved in a variety of molecular processes, including the epigenetic regulation of gene expression, formation of nonmembranous cellular bodies, and sequestration of miRNAs or RNA-binding proteins (RBPs) (reviewed in Kopp and Mendell, 2018; Quinn and Chang, 2016; Wu et al., 2017). While the number of lncRNA genes (15,877) is almost comparable to the number of protein-coding genes (19,881) according to the most recent statistics of human gene annotation (GENCODE GRCh38, http://www.gencodegenes.org/stats.html), their functional annotations are far from complete. For a vast majority of lncRNAs identified to date, our knowledge is limited to their expression patterns and primary sequences (reviewed in de Hoon et al., 2015). Considering that certain fractions of lncRNAs might be “transcriptional noise” produced via stochastic associations of RNA polymerase with open chromatin regions (reviewed in Struhl, 2007), it is necessary to distinguish physiologically relevant lncRNAs from “junk” RNAs. While recent studies identified nuclear-localizing elements in lncRNAs (Lubelsky and Ulitsky, 2018; Zhang et al., 2014), lncRNAs commonly lack conserved sequence motifs or secondary structures, making functional classification of novel lncRNAs rather challenging. This situation is largely different from the case of proteins, which can be systematically categorized into families of molecules according to their individual domain structures (reviewed in Hirose and Nakagawa, 2016).

Recently, a genome-wide CRISPR-interference screen identified hundreds of lncRNAs that affect cellular growth (Liu et al., 2017). Another more specific, genome-scale CRISPR–Cas9 activation screening identified 11 lncRNAs that confer drug resistance in tumor cells (Joung et al., 2017). Further functional lncRNAs might be identified when we can establish appropriate cell-based scalable assays that correctly evaluate the biological functions of lncRNAs in cultured cell lines. On the other hand, lncRNAs tend to display more tissue-and cell-type specific expression patterns (Cabili et al., 2011; Derrien et al., 2012), and their cellular and physiological functions can possibly be addressed only under particular conditions and in certain contexts, some of which might be hard to replicate using commonly available cultured cell lines. In addition, the phenotypes of certain lncRNAs obtained by cellular studies do not always correlate with the phenotypes observed in mutant animals (reviewed in Nakagawa, 2016). Thus, it is essential to perform detailed physiological analyses focusing on specific lncRNAs using mutant animals to understand the biological roles of lncRNAs (reviewed in Bassett et al., 2014; Trapnell et al., 2013). Although recent advancements in genome-editing technologies have enabled rapid generation of genetically modified animals and accelerated reverse-genetic approaches (reviewed in Burgio, 2018), the number of lncRNAs expressed from the genome far exceeds the number of genes that can be feasibly studied using mutant animals. Accordingly, it is important to develop a method that can efficiently identify candidate lncRNAs for subsequent extensive physiological analyses.

Most of the functional lncRNAs examined to date exert their functions through direct association with multiple proteins. Xist, a key factor that controls X chromosome inactivation, associates with a number of RBPs, including hnRNP U, hnRNP K, and Rbm15, as well as transcriptional regulators such as Spen (Chu et al., 2015; Hasegawa et al., 2010; McHugh et al., 2015; Monfort et al., 2015). A number of lncRNAs have also been demonstrated to associate with chromatin-modifying complexes to regulate epigenetic gene expression (reviewed in Holoch and Moazed, 2015). Neat1 functions as an architectural component of nuclear bodies called paraspeckles (Chen and Carmichael, 2009; Clemson et al., 2009; Sasaki et al., 2009; Sunwoo et al., 2009) and builds a characteristic core-shell structure through association with DBHS family members of RBPs and a spectrum of RBPs with prion-like domains (PrLDs), including Fus, Tardbp, and Rbm14 (Fox et al., 2002; Hennig et al., 2015; Naganuma and Hirose, 2013; Souquere et al., 2010; West et al., 2016). These two nuclear lncRNAs, as well as Malat1 and Gomafu/MIAT, which associate with splicing factors (Tano et al., 2010; Tripathi et al., 2010; Tsuiji et al., 2011) form extremely large RNA-protein complexes and are sedimented into fractions heavier than polysomes by sucrose density gradient ultracentrifugation (Ishizuka et al., 2014). NORAD, a cytoplasmic lncRNA that controls genome stability, also associates with an RBP, Pumilio through tandem arrays of its binding motif (Lee et al., 2016; Tichon et al., 2016). These observations led us to hypothesize that the association of multiple proteins might be regarded as a hallmark of functional lncRNAs that exert their functions as ribonucleoprotein complexes.

Some time ago, a simple method termed FAIRE-Seq was developed and enabled the identification of nucleosome-free regions of the genome associated with regulatory proteins (Giresi et al., 2007). FAIRE-Seq utilizes the differential biochemical properties of nucleic acid and proteins during a phenol-chloroform extraction: DNA fragments crosslinked to proteins are separated into the interphase that contains denatured proteins, whereas free DNA is separated into the aqueous phase and is efficiently recovered by ethanol precipitation for subsequent analysis by deep sequencing (Giresi et al., 2007). The success of this method encouraged us to take a similar approach to develop a method that enables the prediction of the functionality of lncRNAs based on differential sensitivity to UV irradiation, which induces covalent bonds between nucleic acids and closely associated proteins (Greenberg, 1979; Wagenmakers et al., 1980). In this study, we irradiated cells with UV and performed RNA-Seq analyses using RNAs purified by conventional acid guanidinium phenol-chloroform extraction (UPA-Seq: <u>U</u>V-<u>p</u>henol <u>a</u>queous phase RNA sequencing). We found that known functional lncRNAs including Xist, Neat1, Malat1, and Gomafu/Miat were efficiently crosslinked to proteins and were largely depleted from the aqueous phase upon UV irradiation, leading to a dramatic decrease in the number of mapped reads obtained by UPA-Seq. On the other hand, the number of reads mapped to Blustr and Upperhand, for which transcription but not transcribed products are important for function (Anderson et al., 2016; Engreitz et al., 2016), were not affected by UV irradiation. We also examined the subcellular distribution of novel functional lncRNA candidates identified by the decrease in UPA-Seq reads and found that some of them formed a characteristic “cloud” at their putative transcription sites. We propose UPA-Seq as a useful tool to select candidate lncRNAs prior to intensive functional analyses.

## Results

### Recovery of known representative functional lncRNAs from the aqueous phase is greatly reduced upon UV irradiation

To investigate whether lncRNAs tightly associate with multiple RBPs, we focused on UV irradiation, which induces the formation of covalent bonds between RNAs and RBPs at low efficiency (Greenberg, 1979; Wagenmakers et al., 1980). Because of the low efficiency of UV crosslinking, we speculated that lncRNAs that weakly associate with interacting proteins remain uncrosslinked under mild UV irradiation conditions. However, functional lncRNAs that form tight ribonucleoprotein complexes are expected to be crosslinked to at least one of the associating proteins, which are then co-fractionated into the interphase after phenol-chloroform extraction (**Fig. 1A**). To test these ideas, we first irradiated the mouse neuroblastoma cell line Neuro2a expressing Gomafu (Sone et al., 2007) with UV_260_ (120 mJ/cm^2^) and examined the amount of RNA recovered from the aqueous phase after conventional acid guanidinium thiocyanate-phenol-chloroform (AGTPC) extraction. The total amount of RNA obtained from UV-irradiated and nonirradiated Neuro2a cells (1 × 10^6^ cells) was 371±11 μg and 264±23 μg, respectively (**Fig. 1B**), suggesting that 29% of total RNA was crosslinked to certain proteins and failed to be recovered from the aqueous phase. Subsequent RT-qPCR analyses revealed that lncRNAs such as Xist, Gomafu, and Malat1 were dramatically reduced upon UV irradiation, whereas the mRNA of a housekeeping gene Gapdh was much less affected (**Fig. 1C**). Interestingly, ribosomal RNAs were also less affected than known functional lncRNAs (**Fig. 1C**), which was consistent with a classic study showing that ribosomal RNAs are less efficiently crosslinked to proteins than other heterologous RNAs (Wagenmakers et al., 1980).

**Figure 1.**
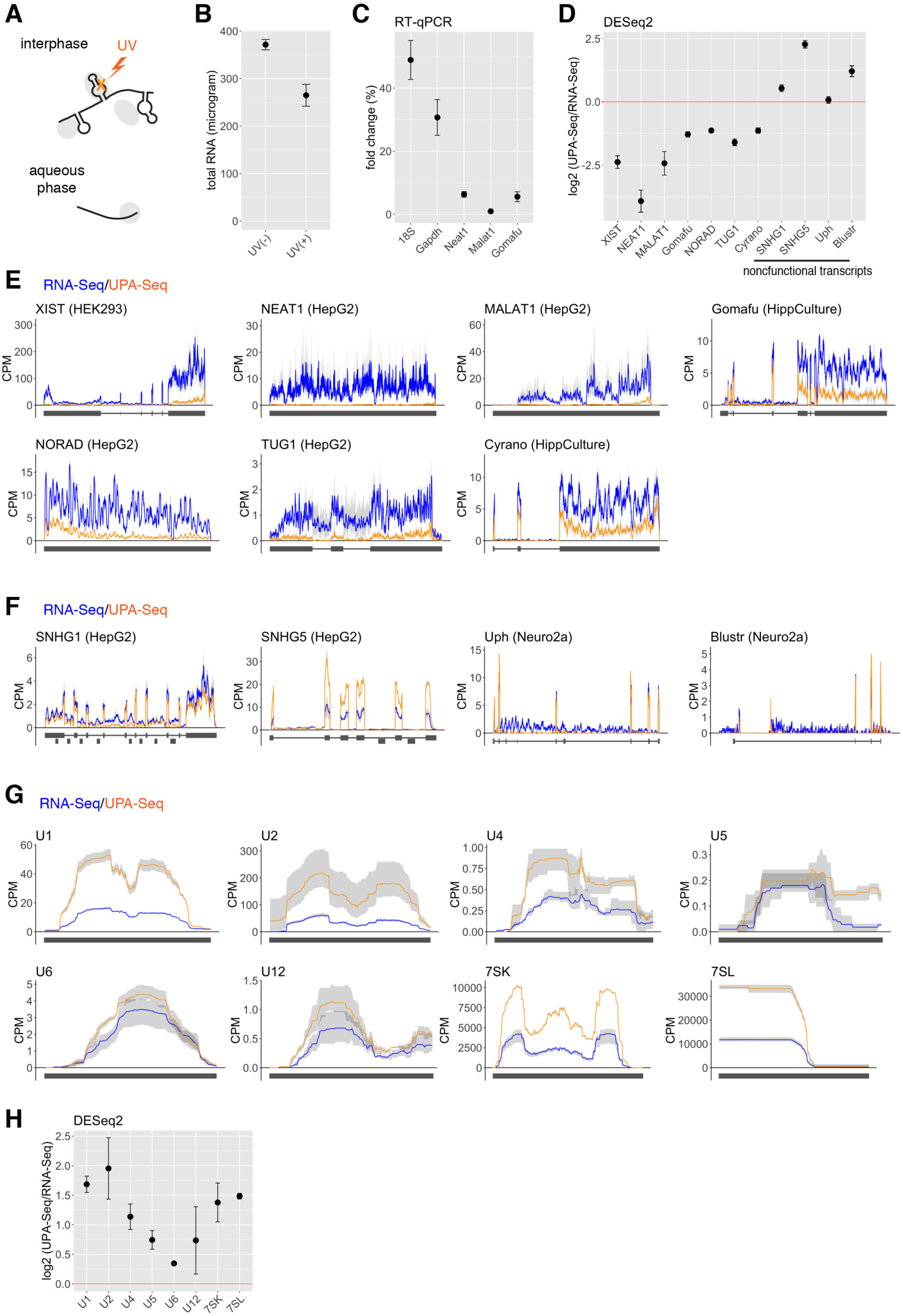
Reduced recovery of known functional lncRNAs from the aqueous phase upon UV irradiation. (A) Schematic drawings of the principle of UPA-Seq. UV irradiation introduces crosslinking between RNA (black lines) and proteins (gray ovals), and crosslinked complexes are separated into the interphase after phenol-chloroform extraction. (B) The amount of total RNA recovered from the aqueous phase before (UV (-)) and after (UV (+))UV irradiation. Data are presented as the means ± s.d. of 3 biological triplicates. 18S: 18S ribosomal RNA. (C) RT-qPCR analysis of the fold change (%) of RNA transcripts recovered from the aqueous phase upon UV irradiation. Note that the recovery of representative functional lncRNA (Neat1, Malat1, and Gomafu) is dramatically decreased upon UV irradiation. Data are presented as the mean ± s.d. of 3 biological triplicates. (D) Log_2_ fold change of reads obtained by UPA-Seq and conventional RNA-Seq mapped to lncRNAs. Sequence reads were counted by featureCount and normalized by DESeq2. Note that nonfunctional lncRNA transcripts do not exhibit decreased recovery. (E) Distribution of UPA-Seq and RNA-Seq reads mapped to representative functional lncRNAs. Blue and orange lines represent the distribution of reads obtained by UPA-Seq and RNA-Seq, respectively. Cell types used for the analysis are indicated by the name of the lncRNA genes. (F) Distribution of UPA-Seq and RNA-Seq reads mapped to lncRNAs, of which transcribed products are thought to be nonfunctional. (G) Distribution of UPA-Seq and RNA-Seq reads mapped to classic ncRNAs. (H) Log_2_ fold change of reads mapped to classic ncRNAs upon UV irradiation. Data are presented as the means ± s.d. of 3 biological triplicates. CPM in E-G represents counts per millions of reads.

To further investigate the sensitivity of lncRNAs to UV irradiation systematically, we performed deep sequencing analysis of the RNAs recovered from the aqueous phase after UV irradiation (UPA-Seq) using multiple cultured cell, including Neuro2a, the human hepatocellular carcinoma cell line HepG2, and the human embryonic kidney cell line HEK293, as well as primary cultures of mouse hippocampal neurons (HippCulture). As a control, we performed conventional RNA sequencing (RNA-Seq) using nonirradiated cells. Comparison of the sequence reads obtained by RNA-Seq and UPA-Seq revealed that the number of sequence reads mapped to representative functional lncRNAs, including Xist, Neat1, Malat1, Gomafu, NORAD, TUG1, and Cyrano, were greatly reduced after UV irradiation (**Fig. 1D, E**). On the other hand, the UPA-Seq reads that mapped to nonfunctional lncRNAs, such as host genes for snoRNA, SNHG1 and SNHG4, were unchanged or rather increased (**Fig. 1D, F**). In addition, UPA-Seq reads that mapped to the exonic regions of Blustr and Upperhand (Uph) were also unchanged (**Fig. 1D, F**), consistent with a previous proposal that the transcription of these lncRNAs, but not their transcribed RNA products, are essential for their biological functions (Anderson et al., 2016; Engreitz et al., 2016). We also examined the fold change of reads that mapped to classic noncoding RNAs associated with the housekeeping process of gene expression, including UsnRNAs, 7SK, and 7SL (**Fig. 1G, H**). Unexpectedly, the sequence reads obtained by UPA-Seq were unchanged or even increased compared to those obtained by conventional RNA-Seq in all of the cases (**Fig. 1G, H**), suggesting that these “functional” noncoding RNAs are less likely to be UV-crosslinked to associating proteins under the irradiation condition, as was the case for ribosomal RNAs previously reported (Wagenmakers et al., 1980).

### Fewer sequence reads are mapped to lncRNAs in UPA-Seq

We then examined the genome-wide fold change of reads mapped to lncRNAs and mRNAs upon UV irradiation (**Fig. 2A**). For this analysis, lncRNAs were defined when they had the GENCODE gene/transcript type annotations “lincRNA”, “antisense”, “bidirectional_promoter_lncRNA”, or “3prime_overlapping_ncRNA”. As expected, the number of UPA-Seq reads mapped to representative functional lncRNAs (Xist, Neat1, Malat1, Gomafu, NORAD, TUG1) was remarkably decreased in all of the four cell types we examined (dark green dots in **Fig. 2A**), providing a rationale to select novel functional lncRNA candidates by the fold change of reads after UV irradiation. We also examined the overall fold change of reads mapped to different GENCODE gene categories upon UV irradiation (**Fig. 2B**). The number of UPA-Seq reads mapped to lncRNAs annotated as “lincRNA” or “antisense” exhibited a significant decrease relative to the counts mapped to protein-coding mRNAs, suggesting that lncRNAs are generally associated with more protein than mRNAs. Notably, the fold change of reads mapping to lncRNAs in HepG2 cells exhibited a bimodal distribution with two peaks, one at approximately zero and the other at approximately-2.6 (**Fig. 2B**), which might represent a group of nonfunctional and functional lncRNAs, respectively. The bimodal distribution of the fold change was also recognizable in Neuro2a cells but not in HEK293 or HippCulture cells, possibly reflecting a differential sensitivity to UV irradiation in each cell type.

**Figure 2.**
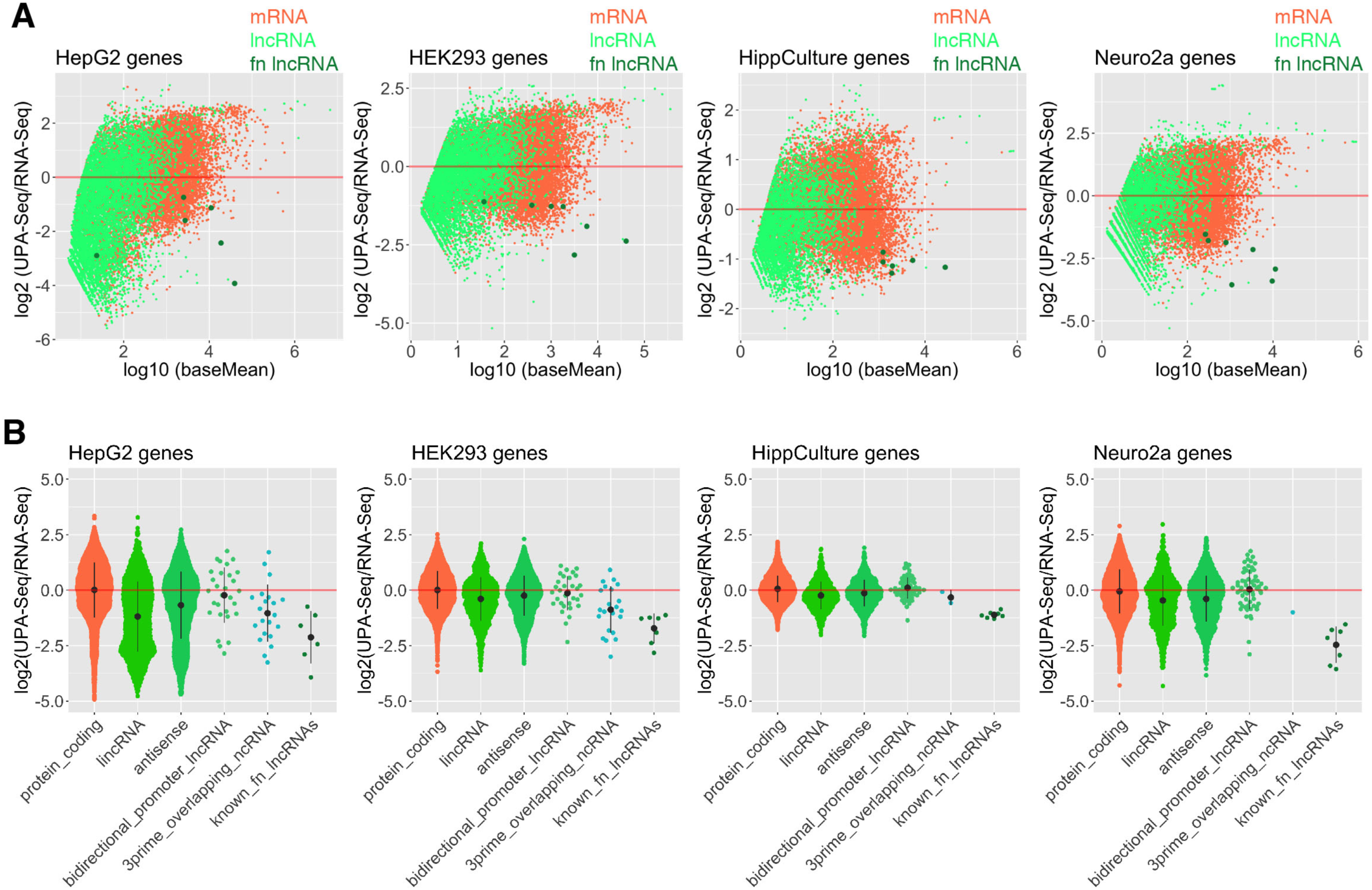
Decreased RNA-Seq reads mapped to known functional lncRNAs upon UV irradiation. (A) MA-plot of log_2_ fold change of reads upon UV irradiation as function of log_10_ baseMeans calculated by DESeq2 in HepG2, HEK293, Neuro2a, and HippCulture cells. Light green dots represent lnRNAs, coral dots represent mRNAs, and dark green dots represent known functional lncRNAs (Xist, Neat1, Malat1, Gomafu, NORAD, TUG1, and Cyrano). Some of the functional lncRNAs that are not expressed in each cell type are not shown in the panels. fn lncRNA: functional lncRNA. (B) Quasirandom beeswarm plots of log_2_ fold change of reads upon UV irradiation in HepG2, HEK293, Neuro2a, and HippCulture cells. Bar plots indicate the means ± s.d. of 3 biological triplicates. Note that lncRNAs generally exhibit negative fold change. known_fn_lncRNAs: known functional lncRNAs.

### Weak correlations between the length or expression levels of lncRNAs and the decrease in UPA-Seq read counts

Because all of the representative functional lncRNAs mentioned above are relatively long and abundantly expressed, the decrease in UPA-Seq reads might be simply explained by increased stochastic interactions between proteins and long, abundant lncRNAs. To test whether this is the case or not, we compared the length and abundance of each lncRNA transcript with the fold change of reads obtained by UPA-Seq and RNA-Seq in HepG2 cells (**Fig. 3A, B**). The Spearman’s rank correlation coefficient (ρ) between the length and the fold change of reads in HepG2 cells was −0.0017 (**Fig. 3A**), suggesting that the decrease in UPA-Seq reads was independent of the length of lncRNAs. We also failed to detect a correlation between the fold change of reads and the expression level of lncRNAs estimated by Reads Per Kilobase of transcript per Million mapped reads (RPKM) (ρ=0.001, **Fig. 3B**). The same trend was observed for all the cell types we used (HEK293, Neuro2a and HippCulture) (**Fig. 3A, B**). These observations suggested that the sensitivity to UV crosslinking is controlled by the amount of RBPs that recognize specific sequences or the secondary structures of lncRNAs rather than by sequence-independent stochastic association with surrounding proteins.

**Figure 3.**
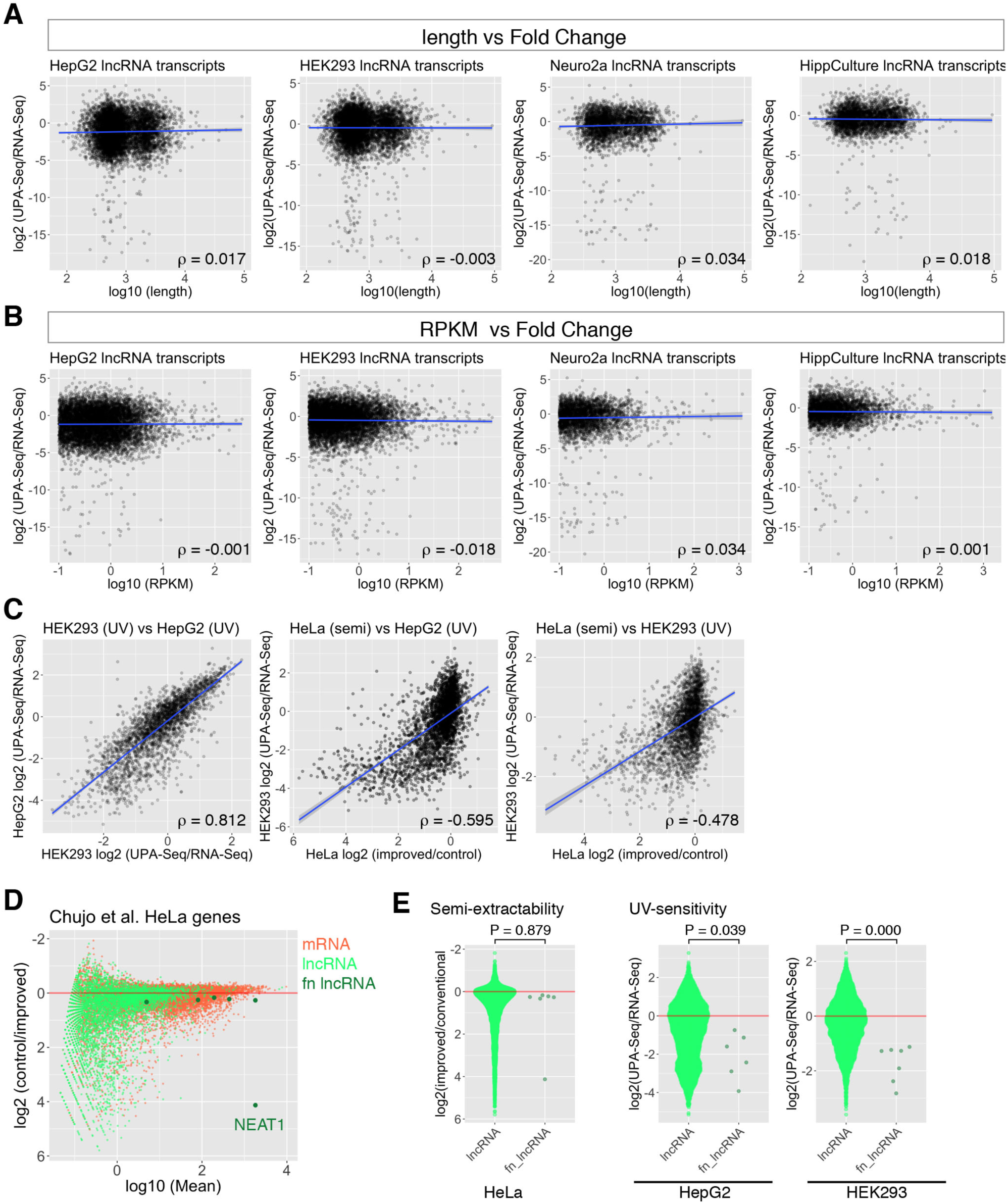
Correlation analysis of fold change of reads upon UV irradiation and the length, RPKM, and semi-extractive properties of lncRNAs. (A, B) Scatter plots illustrating the relationship between the length (A) or RPKM (B) of lncRNAs and the fold change of reads upon UV irradiation in HepG2, HEK293, Neuro2a, and HippCulture cells. Correlation coefficient (ρ) was calculated by Spearman’s rank correlation analyses. Note that the length or RPKM did not exhibit a correlation with the fold change of reads upon UV irradiation. Blue lines represent regression lines. (C) Scatter plots illustrating the relationship between the fold change of reads in two different cell types (HEK293 (UV), HepG2 (UV)) and the fold change of reads after improved extraction (HeLa (semi)) compared with the fold change of reads upon UV irradiation (HepG2 (UV), HEK293 (UV)). Note that the y-axes HeLa (semi) showing the fold change of reads after improved extraction are reversed. Correlation coefficient (ρ) was calculated by Spearman’s rank correlation analyses. Blue lines represent regression lines. (D) MA-plot of log_2_ fold change of reads upon improved extraction as a function of log_10_ baseMeans calculated by DESeq2 with an inverted y-axis. Light green dots represent lnRNAs, coral dots represent mRNAs, and dark green dots represent known functional lncRNA genes (fn lncRNA). Note that the extraction of NEAT1 is dramatically enhanced by the improved extraction, whereas other representative functional lncRNAs exhibited only mild, if any, enhancement. (E) Quasirandom beeswarm plots of log_2_ fold change of reads upon improved extraction (Semi-extractability) and UV irradiation (UV-sensitivity). Cell types used are indicated at the bottom of the plot. P values are calculated by Wilcoxon rank sum test. Note that the y-axis for the semi-extractability in HeLa cell is reversed.

Recently, Chujo et al. discovered that lncRNAs, including NEAT1, that form certain nuclear bodies are hard to purify by conventional AGTPC treatment, and they collectively called them as “semi-extractable” lncRNAs (Chujo et al., 2017). Improved extraction methods such as needle sharing or heat treatment dramatically increased the recovery of semi-extractable lncRNAs, and this semi-extractability has been proposed as a characteristic of architectural lncRNAs that function in nuclear bodies (Chujo et al., 2017). We were interested how these semi-extractable lncRNAs were represented in UPA-Seq read counts, and we compared our results with the dataset reported in the previous study (Chujo et al., 2017). Because the authors used the HeLa cell line, which was different from the cell lines we used, we initially compared the fold change of reads obtained from HepG2 cells and HEK293 cells in our study. Despite the difference in the cell line, we observed a strong correlation between the two values (**Fig. 3C** left panel, ρ=0.812 in Spearman’s rank correlation analyses), suggesting that the fold change of reads mapped to certain lncRNAs upon UV irradiation is fairly conserved across different cell lines. We then compared the fold change of reads with an improved extraction method and the fold change of reads upon UV irradiation (**Fig. 3C** middle and right panels). We observed a fairly good negative correlation between the two values (ρ =-0.595 in HepG2 cells and ρ =-0.4878 in HEK293 cells), suggesting that the decrease in the read counts in UPA-Seq could also be used to predict semi-extractable, architectural lncRNAs. We note that this observation may even be underestimated because of the use of different cell lines for each approach. Notably, the recovery of many of the representative functional lncRNAs, including MALAT1, NORAD, CYRANO, TUG1 and GOMAFU, were not dramatically affected by the improved extraction, whereas they exhibited a remarkable reduction in UPA-Seq (**Fig. 3D, E**). Accordingly, compared to previous methods, UPA-Seq may be applicable for the prediction of a wider range of functional lncRNAs.

### Nuclear localization of functional lncRNA candidates identified by UPA-Seq

We then attempted to identify novel functional lncRNA candidates using UPA-Seq, initially focusing on genes expressed in HippCulture cells. We selected 242 moderately abundant lncRNAs that exhibited decreased read counts in UPA-Seq (log_10_ (baseMean) >1.5, FDR < 0.05) in HippCulture cells (**Fig. 4A**). The majority (86%) of these lncRNAs with GENCODE assigned gene names containing the prefix RP, Gm, or AC or the suffix Rik have not been described in the literature (**Fig. 4B**). We then arbitrarily selected 8 genes from these unannotated lncRNAs as well as 2 genes with annotated names that were not well studied and examined their subcellular distribution using fluorescent in situ hybridization (FISH) (**Fig. 4C**). Typically, we observed one or two discrete dots in the nucleus, and diffuse signals in nucleoplasm were also observed with probes that detect RP24-122I22.2 or RP23-316B4.2 (**Fig. 4C**). We also examined the subcellular distribution of three of the functional lncRNA candidates selected from HepG2 cells (**Fig. 4A, D**). To confirm the discrete dots observed for some of the lncRNAs corresponded to their transcription sites, we focused on RP11-113K21.5, which is located upstream of Rab30 and is conserved between humans and mouse (AC104921.1 in mouse) (**Fig. 4E**). To identify the site of transcription, we performed FISH using a probe that detects an intron of Rab30, a gene located immediately upstream of RP11-113K21.5, which should visualize the genomic locus (**Fig. 4E**). Transcripts of RP11-113K21.5 were observed as two discrete dots in HepG2 cells, which were closely associated with the signals detected by the Rab30 intron probe (**Fig. 4E**). These observations implied that RP11-113K21.5 accumulated at the site of transcription and controlled the expression of neighboring genes, as described for novel lncRNAs identified in a previous study (Joung et al., 2017).

**Figure 4.**
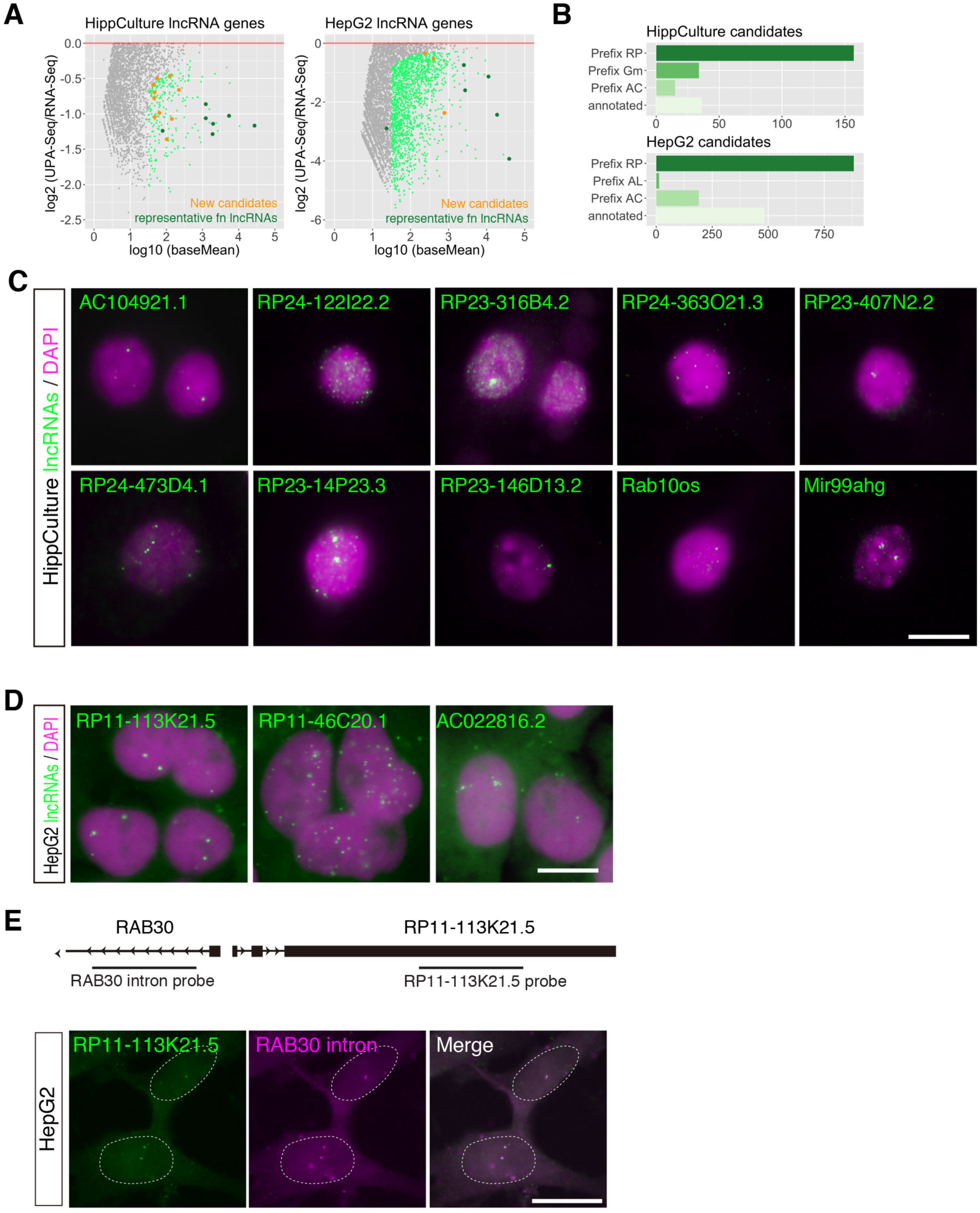
Identification of novel functional lncRNA candidates by UPA-Seq. (A) MA-plot of log_2_ fold change of reads upon UV irradiation in HippCulture and HepG2 cells as a function of log_10_ baseMeans. Light green dots represent lncRNAs that exhibited significant (FDR < 0.05) fold change, orange dots represent functional lncRNA candidates used for the following FISH studies, and dark green dots represent known functional lncRNAs. Note that only lncRNAs with negative log_2_ fold change values are illustrated. (B) Group of functional lncRNA candidates that exhibited significant decrease (FDR < 0.05) upon UV irradiation. Genes are categorized into known genes (annotated) and unknown genes assigned with ID numbers with various prefixes (RP, Gm, AC, and AL). (C, D) FISH images showing the subcellular distribution of functional lncRNA candidates (green) in HippCulture cells (C) and HepG2 cells (D). Magenta denotes nuclei counterstained with DAPI. Scale bar, 10 μm. (E) Schematic illustration showing the gene organization at the Rab30/RP11-113K21.5 locus and the positions of the probes used for simultaneous detection. Note that the signals obtained with probes that detect RP11-113K21.5 were closely associated with the signals obtained with probes that detect Rab30 intron sequences. Dashed lines represent the position of the nucleus. Scale Bar, 10 μm.

### LncRNAs that associate with the prion-like domain-containing RBPs exhibit larger decreases in UPA-Seq reads

All of the results described above suggested that functional lncRNA were easily crosslinked to associated proteins and that their recovery from the aqueous phase was remarkably decreased upon UV irradiation. To investigate the types of RBPs that preferentially associate with functional lncRNAs, we reanalyzed the ENCODE eCLIP resources (https://www.encodeproject.org/eclip/) in which the binding sites of various RBPs were systematically studied (Van Nostrand et al., 2016). We initially calculated the sum of eCLIP enrichment values (number of eCLIP reads/number of size-matched input control reads) for each protein, assuming that the values should reflect certain aspects of protein-binding at least in part. Unexpectedly, we observed a mild positive correlation between the eCLIP enrichment and the fold change of reads upon UV irradiation (**Fig. 5A**), suggesting that association with proteins in general did not necessarily result in decreased read counts in UPA-Seq. We also examined the cumulative distribution of lncRNAs classified according to the total eCLIP enrichment value and confirmed that lncRNAs with lower eCLIP enrichment values were represented in a group of lncRNAs that exhibited greater decreases in read counts in UPA-Seq (**Fig. 5B**). These unexpected results led us to speculate that each RBP differentially contributed to the decrease in UPA-Seq reads and thus examined the cumulative distribution of target lncRNAs for each RBP along with the fold change of reads after UV irradiation. As expected, lncRNAs targeted by each RBP were differentially represented along with the fold change of reads obtained by UPA-Seq (**Fig. 5C, D**). Interestingly, RBPs that contain PrLDs (or an intrinsically disordered domain) were frequently represented in a group of RBPs that associated with lncRNAs exhibiting a decreased recovery from the aqueous phase upon UV irradiation (asterisks in **Fig. 5D**). Comparison of the fold change of reads mapped to target lncRNAs at the median revealed that the association of PrLD-containing RBPs significantly reduced the recovery of lncRNAs upon UV irradiation compared with the association of other RBPs (P=0.037 in Wilcoxon rank sum test) (**Fig. 5E**).

**Figure 5.**
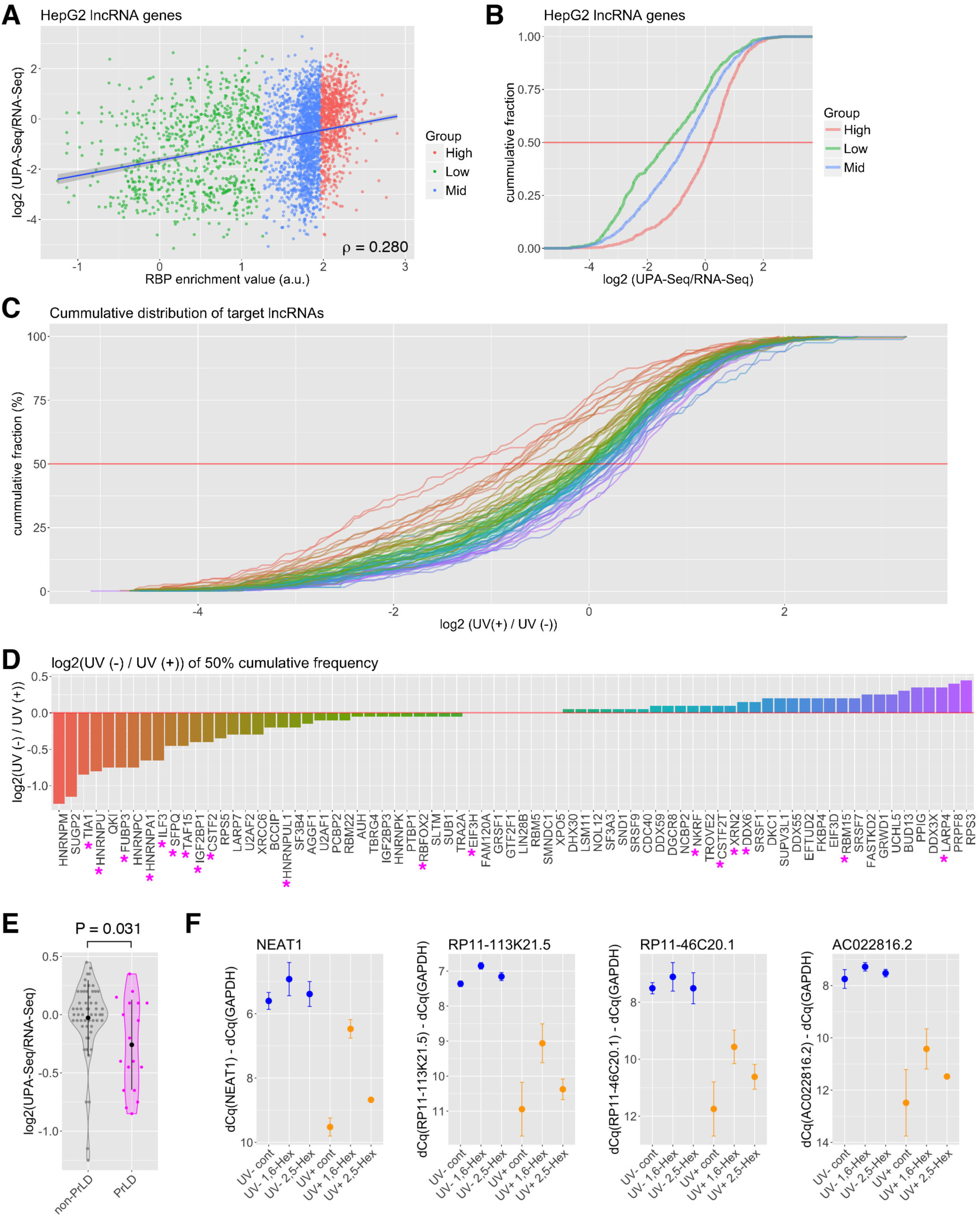
Association with PrLD-containing RBPs preferentially decreases UPA-Seq reads. (A) Scatter plot illustrating the relationship between the total RBP enrichment values (sum of eCLIP/control input reads) and the fold change of reads upon UV irradiation. Blue line represents the regression line. (B) Cumulative distribution of lncRNAs that possessed high (top 25% quintile, red), middle (25-75 percentile, blue), and low (bottom 25% quintile, green) total RBP enrichment values along the fold change of reads upon UV irradiation. Note that lncRNAs with higher total enrichment values tend to be represented in a group that are efficiently recovered from the aqueous phase upon UV irradiation. (C) Cumulative distribution of lncRNAs bound by a series of RBPs shown in D along with the fold change of reads upon UV irradiation. Note that each RBP contributes differentially to the decreased UPA-Seq reads upon UV irradiation. (D) Bar plots illustrating the fold change of reads mapped to lncRNA at the median targeted by indicated RBPs. The magenta asterisks represent RBPs containing a PrLD. (E) Violin and quasirandom beeswarm plots showing the fold change of reads mapped to lncRNA at the median targeted by RBPs without PrLD (non-PrLD) and with PrLD (PrLD). Bar plots indicate the mean ± s.d. of fold change values shown in each category. (F) RT-qPCR quantification of specific lncRNAs indicated at the top of each panel from UV-irradiated (UV+) and nonirradiated (UV-) cells pretreated with control DMSO (cont), 1,6-hexanediol (1,6-Hex), and 2,5-hexanediol (2,5-Hex). The 2,5-hexanediol treatment suppressed the reduction of recovery from the aqueous phase upon UV irradiation. The reversed Y-axes represent the delta Cq values of each lncRNA relative to GAPDH. Bar plots indicate the mean ± s.d. of 3 biological triplicates.

A series of recent studies demonstrated that PrLD-containing RBPs induce liquid-liquid phase separation or hydrogel formation in vitro, providing a molecular basis for the formation of nonmembranous cellular bodies in vivo (reviewed in Uversky, 2017), including the paraspeckles assembled on Neat1 (reviewed in Fox et al., 2018). To further test whether the phase separation of PrLD-containing proteins and associated lncRNAs facilitated covalent bond formation upon UV irradiation, we treated HepG2 cells with 1,6-hexanediol, an aliphatic alcohol that disrupts weak hydrophobic interactions and interferes with liquid-liquid phase separation (Molliex et al., 2015; Ribbeck and Gorlich, 2002; Strom et al., 2017). We also treated the cells with 2,5-hexanediol, which is less effective in disrupting phase-separated cellular bodies. Subsequent RT-qPCR analyses revealed that hexanediol restored the recovery of NEAT1 as well as the novel functional lncRNA candidates RP11-113K21.5, RP11-46C20.1, and AC022816.2 from the aqueous phase upon UV irradiation (**Fig. 5F**). These observations suggested that functional lncRNAs form tight ribonucleoprotein complexes that are easily crosslinked by UV irradiation through interactions with PrLD-containing proteins.

### Reduction of UPA-Seq reads mapped to localizing mRNAs

Although we observed a distinct reduction in UPA-Seq reads that mapped to functional lncRNAs, a certain portion of protein-coding mRNAs also exhibited a comparable decrease in UPA-Seq reads (**Fig. 2A, B**). To obtain insight into the physiological relevance of this reduction, we compared the correlation between the fold change of reads upon UV irradiation and the length of three regions of mRNAs, the 5’ and 3’ untranslated regions (UTR) and the open reading frame (ORF) (**Fig. 6A-C**). Interestingly, the length of 3’ UTR correlated with the decrease in UPA-Seq reads in all of the cell types we examined (**Fig. 6C, D**), whereas no correlation was observed for the length of5’ UTR (**Fig. 6A, D**). The length of the ORF exhibited only a mild correlation in three of the cell types, and no correlation was observed in HippCulture cells (**Fig. 6B, D**). As was the case for lncRNAs, mRNAs that associated with RBPs containing PrLDs exhibited a greater negative fold change upon UV irradiation (**Fig. 6E, F**). We speculated that mRNAs were efficiently crosslinked to PrLD-containing RBPs when they are localized in phase-separated cellular bodies. To test this idea, we examined the cumulative distribution of mRNAs enriched in P-bodies, representative phase-separated nonmembranous cellular bodies (reviewed in Luo et al., 2018), described in Hubstenberger et al. (Hubstenberger et al., 2017) along with the fold change of reads upon UV irradiation in our dataset. As expected, the P-bodies-enriched mRNAs (log_2_ fold change > 2) were preferentially represented in a group that exhibited a greater negative fold change upon UV irradiation (**Fig. 6G, H**). We also examined the cumulative distribution of dendrite or axon-localizing mRNAs described in Cajigas et al. (Cajigas et al., 2012), many of which are known to localize to neuronal RNA granules (reviewed in Hirokawa, 2006). Again, we observed that these synaptic/dendritic neuronal mRNAs were preferentially represented in a group that exhibited decrease in UPA-Seq reads (**Fig. 6I, J**).

**Figure 6.**
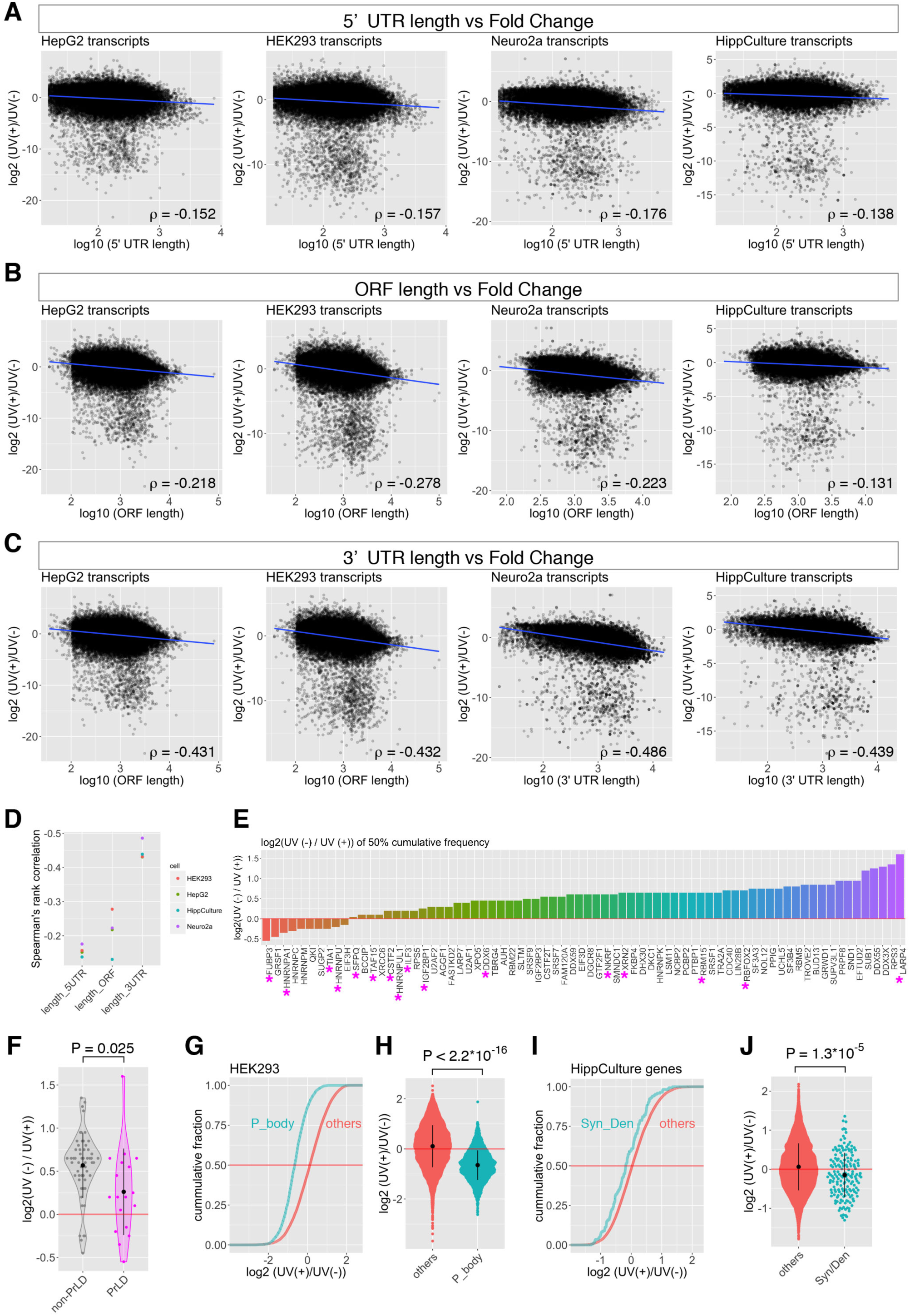
Properties of mRNAs that exhibit reduced recovery from the aqueous phase upon UV irradiation. (A-C) Scatter plots illustrating the relationship between the lengths of 5’ UTR (A), ORF (B), and 3’ UTR (C) of protein-coding mRNAs and the fold change of reads upon UV irradiation. Blue lines indicate the regression line. Correlation coefficient (ρ) values were calculated by Spearman’s rank correlation analysis. (D) Point plots of the Spearman’s rank correlation coefficient ρ value shown in A-C. (E) Bar plots illustrating the fold change of reads mapped to mRNAs at the median targeted by each RBP. The magenta asterisks represents RBPs containing PrLD. (F) Violin and quasirandom beeswarm plots showing the fold change of reads mapped to mRNAs at the median targeted by RBPs without PrLD (non-PrLD) and with PrLD (PrLD). Bar plots indicate the mean ± s.d. of values in each category. (G) Cumulative distribution of mRNAs enriched in P-bodies (P_body) and other mRNAs (others) along with the fold change of reads upon UV irradiation. (H) Quasirandom beeswarm plot showing the fold change of reads mapped to mRNAs that are enriched in P-bodies (P-body) and other mRNAs (other). Note that mRNAs enriched in P-bodies exhibit greater negative fold change compared to others. P value was calculated by Wilcoxon ran sum test. (I) Cumulative distribution of mRNAs localizing at synapse/dendrite (Syn/Den) and other mRNAs (others) along with the fold change of reads upon UV irradiation. (J) Quasirandom beeswarm plot showing the fold change of reads mapped to mRNAs that localize at the synapse/dendrite (Syn/Den) and other mRNAs (other). Note that synapse/dendrite-localizing mRNAs exhibit greater negative fold change compared to others. P value was calculated by Wilcoxon ran sum test.

## Discussion

We have thus demonstrated that comparison of UPA-Seq and RNA-Seq provides valuable information for the selection of candidate lncRNAs to be analyzed in subsequent studies. We recognize that the reduced recovery from the aqueous phase upon UV irradiation is not a definitive feature for “functional” noncoding RNAs because such reduction was not observed with classic functional noncoding RNAs involved in evolutionary conserved processes of gene expression, including rRNA, UsnRNAs, snoRNAs, 7SK, and 7SL, all of which form well-characterized ribonucleoprotein complexes. Considering that covalent bond formation between the RNA and protein should impair the physiological function of these molecular machineries, nucleotide sequences and amino acid residues of these classic ribonucleoprotein complexes might have been selected to decrease the efficiency of UV crosslinking during the course of evolution. We also failed to detect reduction of UPA-Seq reads mapped to lncRNAs, including Blustr and Handup, which regulate the expression of nearby genes by their transcription but not their transcribed products. Our prediction of functionality is thus applicable to a limited group of lncRNAs that tightly associate with proteins to form large ribonucleoprotein complexes. Nevertheless, we propose UPA-Seq should be useful for selecting candidate genes involved in the processes of interest during the initial screening step, due to its technical simplicity and compatibility.

Although all of the cell types we used exhibited similar reduction in UPA-Seq reads, they responded differentially to UV irradiation: the fold change of reads mapped to lncRNAs in HepG2 cells and Neuro2a cells exhibited greater negative values and bimodal distributions, while HEK293 cells and HippCulture cells exhibited a milder decrease and moderate Gaussian distributions. The differential responses might represent differential association of lncRNAs with proteins specifically expressed in each cell type. Alternatively, lncRNAs receive different amounts of UV energy due to cell type-specific composition of molecules that absorb UV light, including various metabolites as well as the residues of proteins and nucleic acids that are differentially expressed. Regardless, the condition of UV irradiation must be carefully optimized for each cell type for maximal use of UPA-Seq to select functional lncRNA candidates.

Importantly, lncRNAs that exhibited greater decreases in UPA-Seq reads preferentially associated with RBPs containing the PrLD. A series of studies revealed that one of the characteristic functions of lncRNAs is to form nonmembranous organelles or a ribonucleoprotein milieu within the cells via liquid-liquid phase separation through interaction with PrLD-containing RBPs (Banani et al., 2017; Fay and Anderson, 2018; Fay et al., 2017; Hennig et al., 2015; Maharana et al., 2018), and UPA-Seq may be useful for identifying novel lncRNAs included in phase-separated cellular bodies. Actually, recently reported semi-extractable lncRNAs that form novel nuclear bodies were represented in a group that exhibited greater decreases in UPA-Seq reads, and we did observe the formation of a distinct RNA “cloud” around the putative transcription sites for RP23-316B4.2 and RP23-14P23.3. Further functional studies on these lncRNA candidates will expand our knowledge of the biological processes regulated by this group of molecules.

While we primarily focused on lncRNAs in this study, certain fractions of mRNAs also exhibited a dramatic decrease in UPA-Seq reads. Interestingly, the length of 3’ UTR exhibited good correlation with fold changes in reads upon UV irradiation, which was consistent with the well-established concept that the subcellular localization of mRNAs is regulated by 3’ UTR regions through their interaction with miscellaneous RBPs (reviewed in Mayr, 2016). We observed preferential representation of P-bodies-enriched and neurite-localizing mRNAs among the genes that exhibited greater decreases in reads upon UV irradiation, which preferentially associated with proteins containing PrLD. UPA-Seq might thus be useful to make a candidate list of mRNAs that are localized in phase-separated nonmembranous cellular bodies including, stress granules, neuronal granules, and germinal granules.

Overall, the simple combination of UV irradiation and phenol-chloroform extraction provides a versatile tool for the prediction of candidate lncRNAs that tightly associate with regulatory proteins. It would be intriguing if we could predict functionality of lncRNAs in the future by combining information obtained by scalable deep sequencing analyses including UPA-Seq, eCLIP, and various structure-probing technologies recently developed (reviewed in Silverman et al., 2016).

## Acknowledgements

We would like to thank Mr. Shaolong Lin and Ms. Aina Takemoto for initial input for the study, Kyoko Chiba for technical advice on hippocampal cultures, Dr. Rei Yoshimoto at Fujita Health University for advice on RNA-Seq data analyses, and members of the RNA Biology Laboratory for active discussion. We also would like to thank the Research Resource Center at RIKEN BSI for technical support. This work used the Vincent J. Coates Genomics Sequencing Laboratory at UC Berkeley, supported by the NIH S10 OD018174 Instrumentation Grant. This work was supported by JSPS KAKENHI Grant Number 17H03604 and MEXT KAKENHI Grant Number 26113005 granted to S.N., JSPS KAKENHI Grant Number 17H04998 and MEXT KAKENHI Grant Number 17H05679IS granted to S.I., and JSPS KAKENHI Grant Number 16H06279 (PAGS).

## Author contributions

TK performed RNA extraction, data analyses, and FISH; KF performed experiments using hexanediol; MM generated libraries for RNA sequencing; SI organized the RNA sequencing; and SN organized the experiments, analyzed data, and wrote the manuscript.

## Experimental Procedures

### Cell culture

HEK293T and Neuro2a cells were cultured in DMEM/Ham’s F-12 (#048-29785, Wako, Japan) supplemented with 10% fetal bovine serum and penicillin-streptomycin. HepG2 cells were cultured in DMEM (#D5796, Sigma Aldrich) supplemented with 10% fetal bovine serum and penicillin-streptomycin. Primary hippocampal neurons were prepared from E16.5 C57/BL6NCr or ICR mouse brain for RNA-Seq and FISH experiments, respectively, following a previously described protocol (Chiba et al., 2014). Briefly, hippocampal tissues were dissected from E16.5 embryos in ice-cold HEPES-buffered saline (HBSS), treated with 2 mg/ml papain (#LSS03119, Worthington) for 20 min at 37°C, and then gently dissociated into single cells by mild pipetting in the presence of 0.2 mg/ml DNase I (#311284932001, Roche). After filtration through a cell strainer (#352350, BD Falcon), cells were then resuspended in culture medium (Neurobasal medium, #21103049, Thermo Fisher) supplemented with B27 (#17504044, Thermo Fisher), L-glutamine and 1% horse serum and then plated over coverslips precoated with poly-L-lysine and polyethylenimine at a density of 3.5 × 10^4^ cells/cm^2^. Three days after the plating, cells were treated with 10 μM AraC for 24 hours to remove the glial cells and then further cultured for 10 days. For hexanediol treatment, HepG2 cells were treated with 15% 1,6-or 2,5-hexanediol (#240117 and H11904, SIGMA) or DMSO for 5 min at room temperature. After treatment, the medium was replaced with HCMF, and the cells were immediately irradiated with UV as described below.

### UV irradiation and RNA extraction

Culture medium was replaced with 1 ml of ice-cold HBSS, and cells were crosslinked at 254 nm UV with 120 mJ/cm^2^ total energy in a Funa UV linker (#FS-800, Funakoshi, Japan) on ice. Total RNA was purified using TRIzol reagent (#15596026, Invitrogen) following the manufacturer’s instructions with an additional heating step at 55°C for 10 min before the addition of 1/5 volume of chloroform.

### RT-qPCR analyses

cDNA was synthesized from 1 μg of total RNA using the ReverTra Ace qPCR RT Master Mix (#FSQ-201, TOYOBO, Japan) following the manufacturer’s instruction. A total of 1/50 of synthesized cDNA and 0.3 μM for each primer were used for the qPCR reactions. RT-qPCR was performed using THUNDERBIRD SYBR qPCR Mix (#QPS-201, TOYOBO) and CFX Connect (BIORAD) with the following conditions: 95°C for 1 min followed by 40 cycles of 95°C for 15 sec and 60°C for 60 sec. qPCR rimers used in this study are listed in supplemental table T1.

### FISH

FISH was performed following a protocol previously described (Mito et al., 2016). Briefly, cells were plated on PLL-coated coverslips and fixed in 4% paraformaldehyde for 10 minutes at room temperature. After permeabilization with 0.5% Triton X-100 in PBS, cells were hybridized with 1 μg/ml DIG-or FITC labeled RNA probes diluted in hybridization buffer for overnight at 55°C. After washing twice with 50% formamide/2x SSC for 30 minutes at 55°C, samples were treated with RNaseA, and further washed with 2X SSC and 0.2X SSC for 30 minutes each at 55°C. Fluorescent images were obtained using an epifluorescence microscope (BX51, Olympus) equipped with a CCD camera (DP70). Probes and antibodies used are described in supplemental table T1.

### RNA-Seq and data analyses

Deep sequencing libraries were made following a standard protocol using Ribo-Zero Gold rRNA Removal Kit (Human/Mouse/Rat) (Illumina) and TruSeq RNA Sample Preparation v2 (Illumina). and deep sequencing was performed using an Illumina Hiseq2500 (Neuro2a, HippCulture, and 293 cells) or Hiseq4000 (HepG2 cells). Sequence reads were mapped to hg38 and mm10 genome builds using TopHat2 (Kim et al., 2013). The numbers of sequence reads mapped to each exon were counted using featureCount (Liao et al., 2014) with-O option (allowMultiOverlap) and analyzed at the gene level. After removing genes that have an average read of 0.5 per millions of reads in UV nonirradiated samples, the data count was analyzed using DESeq2 (Love et al., 2014) to calculate the normalized fold change between the UV-irradiated and nonirradiated samples. For the numbers of reads mapped to each transcript, we used Cuffdiff2 (Trapnell et al., 2013), and low expressers (RPKM < 0.1) were removed for subsequent correlation analyses. The annotations of genes and transcripts was obtained from the GENCODE homepage (https://www.gencodegenes.org/), and GRCh38.p12 and GRCm38.p6 were used for the human and mouse genome annotation, respectively. The lengths of lncRNAs were calculated from the Fasta files downloaded from the GENCODE homepage (long noncoding RNA transcript sequences) using the readDNAstringSet command in the Biostrings package (DOI: 10.18129/B9.bioc.Biostrings). The length of the 5’ UTR, 3’ UTR, and ORF of mRNAs was obtained from the foldUtr5, knownGenePep, and foldUTR3 tables in the GENCODE/UCSC genes track for human/mouse, which was downloaded using the Table Browser of the UCSC genome browser (http://genome.ucsc.edu/index.html). Coverage plots, beeswarm plots, scatter plots, bar plots, cumulative plots, and violin plots were drawn with ggplot2 (Wickham, 2009). For eCLIP analyses, bam files of eCLIP and mock input controls for each RBP were downloaded from the ENCODE homepage (https://www.encodeproject.org/search/?type=Experiment&assay_title=eCLIP), and the numbers of reads mapped to each lncRNA and mRNAs were counted using featureCount with-O option (allowMultiOverlap) and analyzed at the gene level. The percent of input (eCLIP/control) values were calculated for each RBP, and RNA transcripts that exhibited 2× enrichment were determined as target lncRNAs/mRNAs.

### Data availability

The sequencing data obtained in this study were deposited at NCBI GEO [GSE114789]. To review GEO accession GSE114789:

---

**Supplemental table T1:**
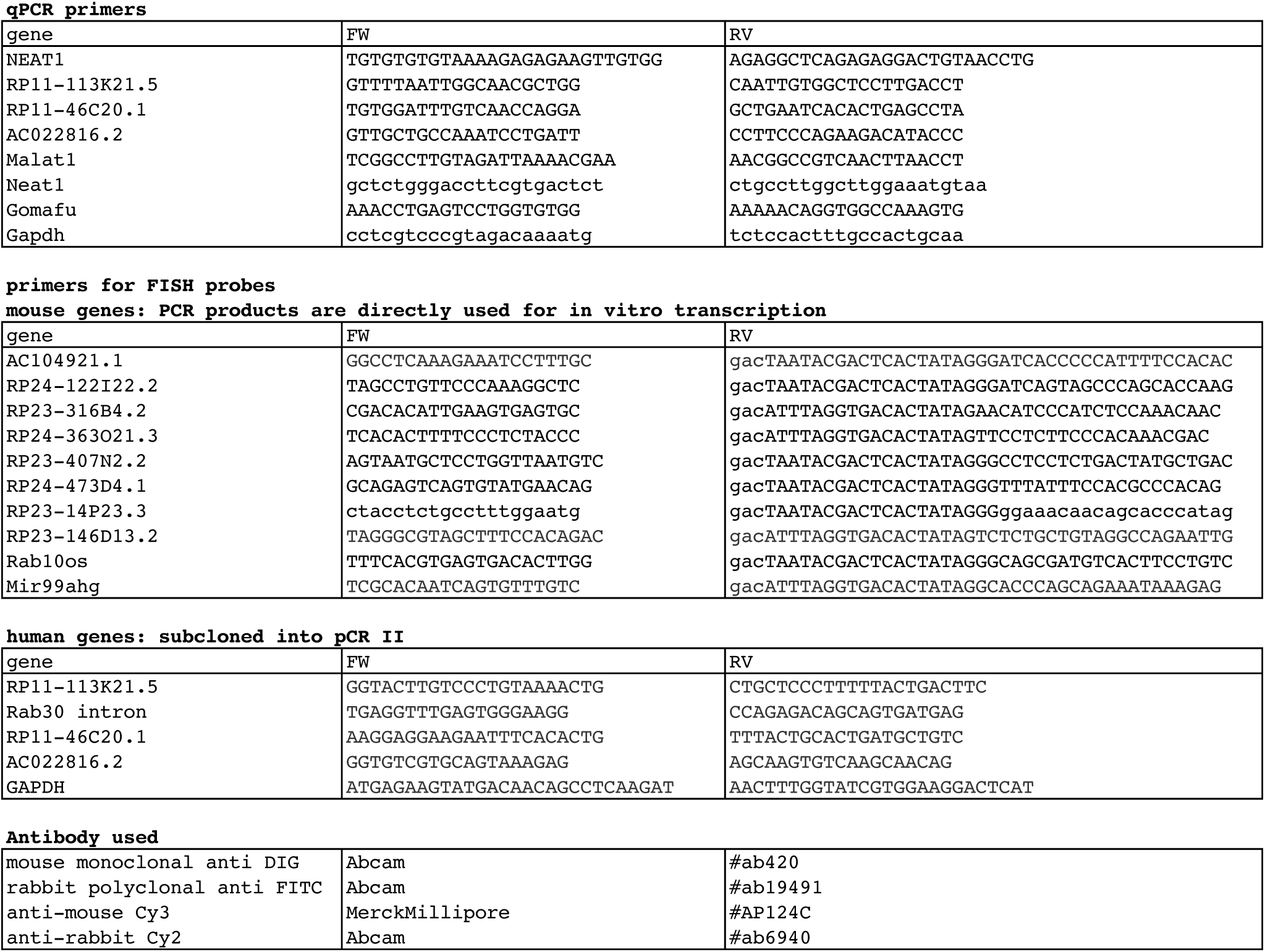
Primers and antibodies used in this study

